# NetTCR-struc, a structure driven approach for prediction of TCR-pMHC interactions

**DOI:** 10.1101/2025.03.22.644721

**Authors:** Sebastian N Deleuran, Morten Nielsen

## Abstract

Accurate modeling of T cell receptor (TCR)–peptide–major histocompatibility complex (pMHC) interactions is critical for understanding immune recognition. In this study, we present advances in structural modeling of TCR-pMHC class I complexes focusing on improving docking quality scoring and structural model selection using graph neural networks (GNN). We find that AlphaFold-Multimer’s confidence score in certain cases correlates poorly with DockQ quality scores, leading to overestimation of model accuracy. Our proposed GNN solution achieves a 25% increase in Spearman’s correlation between predicted quality and DockQ (from 0.681 to 0.855) and improves docking candidate ranking. Additionally, the GNN completely avoids selection of failed structures. Additionally, we assess the ability of our models to distinguish binding from non-binding TCR-pMHC interactions based on their predicted quality. Here, we demonstrate that our proposed model, particularly for high-quality structural models, is capable of discriminating between binding and non-binding complexes in a zero-shot setting. However, our findings also underlined that the structural pipeline struggled to generate sufficiently accurate TCR-pMHC models for reliable binding classification, highlighting the need for further improvements in modeling accuracy.

## Introduction

T cells drive the adaptive immune response by recognizing and eliminating cells displaying foreign peptides through major histocompatibility complexes (MHC) [1]. This process is facilitated by interaction between the T cell receptor (TCR) and the peptide-MHC complex (pMHC) which serves as a crucial checkpoint for immune activation. Understanding the rules governing this interaction is critical, as it is central to the development of a wide range of immunotherapy treatments.

Computational prediction of this event presents an effective avenue of greatly accelerating development of immunotherapies, and consequently a wide range of methods aiming to achieve this have been developed [2], [3], [4], [5], [6]. Primarily, these methods represent the TCR and pMHC using their amino acid sequences, while often utilizing a reduced representation of the TCR by only considering either all CDRs or only the most variable and specificity-defining CDR3 [7], [8]. While some success in the many-shot learning setting has been demonstrated, the zero-shot setting, i.e. inference on completely unseen TCRs and peptides, remains largely unsolved [9]. An estimate of 10^8 unique TCRB sequences may exist in a single individual that, through cross-reactivity and alternative binding modes, may interact with the staggeringly high number of 20^9 9-mer amino acid combinations [10], [11]. Given that approximately only 50,000 paired chain TCR-pHLA class I interactions, not accounting for redundancy, have been described in major databases IEDB and VDJdb, the poor zero-shot performance observed in current state-of-the-art models is unsurprising [2], [12], [13].

Recent advances in protein structure prediction have allowed accurate structural modeling of TCR-pMHC complexes [14], [15], [16].This provides a new avenue in tackling the TCR specificity prediction task, as the conserved nature of protein structure may work as a significantly less diverse perspective on TCRs and peptides.

However, at this time, few methods explicitly utilize structural data and those that do, have not demonstrated significant gains in performance over methods trained on sequence data [17]. Besides the computational cost of performing large scale experiments with state-of-the-art protein modeling tools such as AlphaFold (AF), additional challenges in utilizing structural data are present. Accurate modeling of TCRs, especially in their docked conformations, is an immensely challenging task due to their highly variable and long CDR3 loops. Further analysis of these structural models also remains difficult, given that stimulatory TCR binding can depend on the formation of very few contacts that may not be captured even in high quality models [18]. Recent work by Motmaen et al. and Yin et al. on utilizing AF for modeling pMHC and TCR-pMHC complexes by modifying pMHC and TCR multiple sequence alignment (MSA) and template featurization, have provided significant improvements in modeling accuracy over AlphaFold-M (AF-M) [15], [19]. The latter approach demonstrates high modeling accuracy on a benchmark dataset of solved TCR-pMHC structures. However, the confidence metric provided by AF-M appears to not correlate strongly with the modeling quality of TCR-pMHC interfaces, when quantified with the DockQ metric [20]. Consequently, selection of high quality models from the pool of modeled structures is not guaranteed. Here, we propose a graph neural network based approach for significantly more accurate quality scoring of TCR-pMHC complexes modeled using AF-M. The model is based on geometric vector perceptron layers allowing for fine grained encoding of geometric features, and is trained to perform regression on the DockQ docking quality metric [21]. In order to generate training data for this model, we additionally present results for perturbation of AF MSA and template features and its effect on modeling diversity and quality. Secondly, we apply our docking quality scoring approach to the task of TCR-pMHC binding prediction and demonstrate its efficacy in correctly ranking binding TCR-pMHC complexes when sufficiently accurate structural models are available.

## Material and Methods

### Modeling pipeline

Structures of TCR-pMHC complexes were modeled using an AF-Multimer version 2.3 based pipeline. Multiple sequence alignment (MSA) and template features were created using the approach described by Yin et al. [15]. Here, template features for the pMHC are generated such that the pMHC is modeled as a single chain, allowing for the use of docked pMHC templates. Additionally, TCR MSA and template features are generated from a reduced database of immunoglobulin proteins. In order to increase modeling throughput on high performance compute clusters, the featurization and modeling steps of AF were decoupled so that featurization for a batch of sequences could be run in parallel with modeling once features for the first entry in the batch is completed.

An additional set of options for perturbing MSA and template features were added to the pipeline, with the aim of increasing diversity of the set of structures predicted for a given input [22]. These include random mutation in the MSA, column wise mutation in the MSA, masking of MSA hits (resembling MSA subsampling) and addition of gaussian noise to structural template atomic coordinates. An option to enable dropout of AF modules was also added.

### Training dataset for docking quality scoring

The dataset used for training and evaluating the GVP-GNN DockQ regressor was constructed from a set of solved TCR-pMHC class I complex structures. Structures were obtained from RCSB and TCRs were trimmed to their variable domains [23]. Complexes that contained peptides with non standard amino acids were removed from the dataset. Additionally, structures were filtered to human complexes with α:β TCRs. Finally, a resolution cutoff of 3.5Å was applied resulting in a final data set of 80 structures.

For the cross-validation setup, partitions were created in the following manner. Structures released after the AF-M 2.3 training dataset cutoff of 2021-09-30 were selected for use as a benchmark dataset. The Hobohm 1 algorithm was applied to the data for redundancy reduction, using a 95% sequence similarity threshold. Sequence similarity was calculated over the alignment length, and any complex with a TCRα or TCRβ sequence that was 95% similar to an already encountered sequence was dropped. Following this, for the training data structures, complete linkage agglomerative clustering based on TCRα or TCRβ sequence similarity w as applied in order to generate 5 partitions. Here, the AgglomerativeClustering method from scikit-learn was applied to a matrix of pairwise average TCRα or TCRβ sequence similarities, setting the number of desired clusters to 5 and otherwise using the default settings provided in scikit-learn v1.0.2. The resulting clusters were then used to define the partitions containing 19, 18, 6, 6 and 6 solved structures. The benchmark dataset contained 25 solved structures.

Sequences from the solved structures were extracted and used to generate structural models using the AF-M based pipeline. In order to increase the diversity of modeling quality for each target in the dataset, the input feature perturbation methods described in the Modeling pipeline section were applied on MSA and template features for each modeling seed. Structures were modeled under different pipeline configurations in order to increase the uniformity of the docking pose DockQ distribution. The following runs were performed in order to achieve this:

- No restriction on template selection, 30 candidates per AF model (150 total), and a maximum number of recycling of 3.
- No template information, except for the pMHC, masking of 60% of MSAs, random substitution of 60% of MSA residues, 30 candidates per AF model (150 total).
- No template information, except for the pMHC, masking of 20% of MSAs, random substitution of 20% of MSA residues, 30 candidates per AF mode (150 total).
- Maximum template date set to AF-M 2.3 training dataset cutoff (2021-09-30) and a template and query sequence similarity threshold of 90%, masking of 15% of MSAs, random substitution of 15% of MSA residues, 60 candidates per AF model (300 total).

This pipeline, thus resulted in 750 candidate structural models being generated for each input TCR-pMHC entry. Subsequently models were scored against their ground truth targets using DockQ, considering only the TCR and peptide interactions. As a measure against redundancy, for each target, models from the training/validation partitions were placed in 20 bins according to their DockQ score and up to 20 models from each bin were sampled. Additionally, a second dataset was created where only models with DockQ >= 0.5 were retained, and where no redundancy reduction was performed w.r.t. to the DockQ distribution. Finally, for targets in the benchmark dataset, models with more than 5 TRA-peptide or TRB-peptide backbone clashes were filtered. A backbone clash was defined as backbone atoms in peptide and TCR being within 3Å of each other. This resulted in two training datasets with 12057 (full DockQ range) and 16541 (DockQ >= 0.5 and no homology reduction) models over 55 targets and a benchmark set of 3750 models for 25 targets.

### The GVP-GNN regressor

DockQ regression was performed using a geometric vector perceptron graph neural network (GVP-GNN) [21] (Figure 1). The graph representation of a given protein complex was generated in a per-residue manner, where nodes represented amino acid residues and edges were created based on the euclidean distances of these nodes. As a regularizing measure, the model was trained using a pair mean squared error (MSE) loss function, as described by Jing et al., 2021:

**Figure 1:**
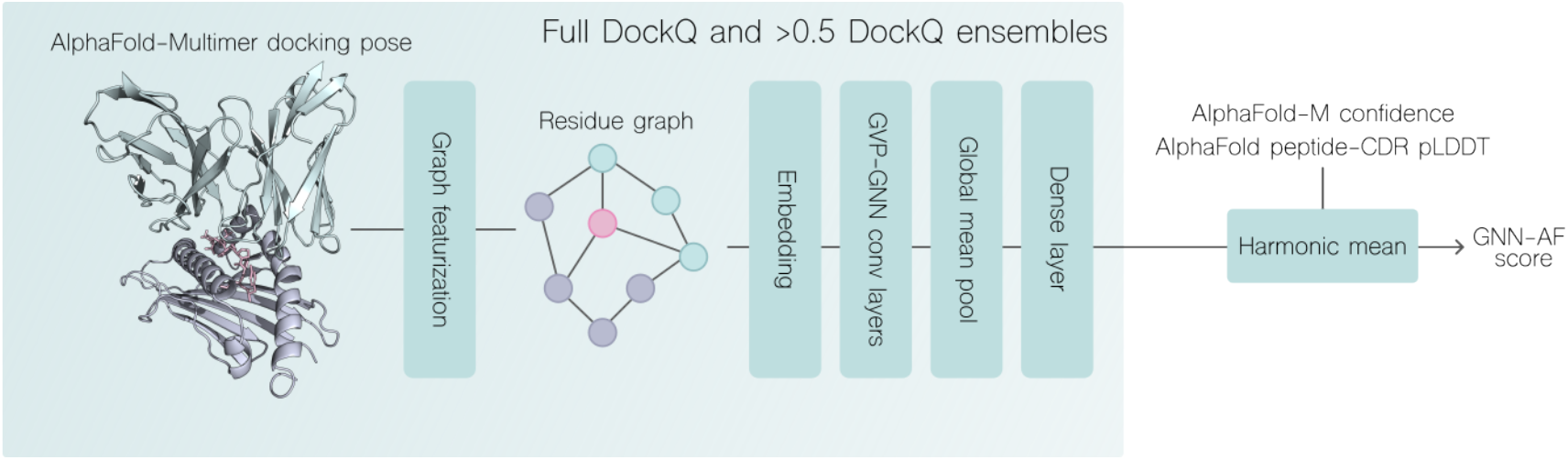
Overview of DockQ regression and quality prediction pipeline. A docking pose generated with AlphaFold-Multimer is featurized as a graph (see materials and methods). Two sets of 5 GNN-GVP models resulting from training on 5 cross-validation splits, where one set was trained only on docking poses with DockQ >0.5 are used to score the structural model. The combined GNN-AF score is computed by the harmonic mean of the two GVP-GNN ensemble scores, AF-M confidence and the peptide-CDR pLDDT score. For more details, see materials and methods.

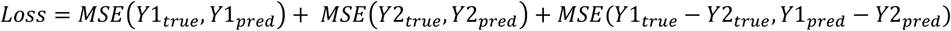

Where Y1_true_, Y1_pred_, Y2_true_, Y2_pred_ are predicted and true DockQ values for the same target structure Y. The difference term, MSE(Y1_true_ -Y2_true_, Y1_pred_ -Y2_pred_), aims to reduce fitting to features that are static between candidates of the same target such as amino acid sequence, by penalizing cases where one candidate is scored more accurately than another candidate, potentially caused by overfitting to these static features.

Models were trained in a 5-fold cross-validation setup to obtain an ensemble of models. We trained two sets of models, one on a dataset composed only of structural models with a DockQ score over 0.5, and one trained on a dataset spanning the full range of the DockQ metric. This resulted in a set of 10 models. Models were constructed and trained with the following hyperparameters:

- node_hidden_dimension_scalar = 50
- node_hidden_dimension_vector = 8
- edge_hidden_dimension_scalar = 50
- node_hidden_dimension_vector = 8
- Graph convolution layers = 3
- dropout rate = 0.5
- optimizer = Adam
- LR scheduler = CosineAnnealingLR
- learning rate = 0.0001
- weight decay = 0.0001
- batch size = 32
- epochs = 100.

We define a GNN ensemble score for a given structure, as the harmonic mean of two predicted DockQ values, which are obtained from taking the arithmetic mean of the 5 GNNs trained on the full DockQ range, and the arithmetic mean of the 5 GNNs trained only on structural models with a >0.5 DockQ.

We additionally define a consensus predicted quality score for a given input as the harmonic mean of the two GNN scores, AF-M confidence metric and CDR-peptide pLDDT. The CDR-peptide pLDDT was computed by extracting the pLDDT scores associated with CDR123αβ and peptide residues, and taking the arithmetic mean of these scores. We term this score GNN-AF. Thus, the predicted quality is given by:

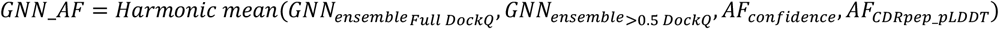

We chose to use the harmonic mean, as it will give more weight to smaller numbers when computing the mean. This is useful, as the AF confidence metric is more prone to overestimation rather than underestimation of quality, and because the graph neural networks developed here can predict DockQ values in the lower range accurately. Thus, when either of these scores are low for a given input, we want to ensure that this input is assigned a low score.

### Data featurization

Graph featurization of protein structures was performed following the methods described in Jing et al., 2021. Here, each node corresponded to a residue in the full protein complex. For each node v_i_, its set of edges was defined as those connecting v_i_ to its 30 nearest neighbors measured by euclidean distance. Scalar and vector node features were defined as suggested by Jing et al., 2021. Let v_i_ be a node representing an amino acid in the i’th position of the concatenated sequence:

- Scalar features residue v_i_:
  - {sin, cos} ° {φ, ψ, ω}, where φ, ψ, ω are the dihedral angles of v_i_.
  - One hot encoding of the amino acid of v_i_.
  - One hot encoding of the chain that v_i_ belongs to.
- Vector features of residue v_i_:
  - Unit vectors describing the relative orientation of v_i-1_ and v_i+1_ w.r.t to their Cα-atoms.
  - Unit vector describing the imputed direction of Cβ_i_ − Cα_i_, computed by:

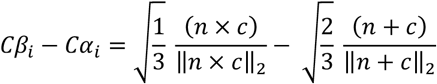

where n = N_i_ − Cα_i_ and c = C_i_ − Cα_i_

Let e_ij_ be a directed edge between two nodes v_i_ and v_j_.

- Scalar features of e_ji_:
  - An encoding of ||Cα_j_ − Cα_i_||^2^ in terms of 16 Gaussian radial basis functions.
  - A sinusoidal positional embedding representing the distance of v_i_ and v_j_ in the primary structure. For inter-chain edges, the positional embeddings were set to 0, as the distance in primary structure is not defined here.
- Vector features of e_ji_:
  - Unit vector describing the direction Cα_j_ − Cα_i_.

We additionally present a GNN model, where we incorporate structural embeddings as additional node scalar features. These embeddings are extracted from the final layer of the encoder module of the ESM-IF1 inverse folding model [24]. Briefly, this model is trained to generate amino acid sequences that are likely to fold into the geometry specified by an input protein structure. The model uses a GVP-GNN module to generate structural features, followed by a generic encoder-decoder transformer module. Each embedding has a shape of 512 x N_AA_ where N_AA_ represents the total length of all chains in a TCR-pMHC complex. For each residue *i*, its corresponding embedding vector was concatenated with the scalar node feature vector for v_i_. We denote this model GNN-ens-IF1, and GNN-IF1 when used in the consensus score.

### TCR specificity data

The binding classification dataset was derived from Jensen (2024) [2]. Briefly, the positive examples in this dataset were derived from IEDB, VDJdb as well as a 10x Genomics sequencing dataset denoised using the ITRAP algorithm [12], [13], [25], [26]. Swapped negatives were generated by mismatching each positive TCRs - pMHC pair with 5 TCRs positive to other pMHCs. In this study, we opted to downsampled the dataset with up to 200 positive examples per peptide and resample negative examples following the same procedure as described by the authors. Partitions were then generated from this downsampled dataset, again using the procedure described by the authors. For more details, we refer to the methods section of Jensen (2024). The downsampled dataset was composed of 2945 binding complexes and 14725 (5 * 2945) swapped complexes giving a total of 17670 examples.

### Modeling of TCR-pMHC binding dataset

For each set of sequences in the retrieved TCR-pMHC binding dataset, 10 candidates for each of the 5 AlphaFold-M multimer models were modeled in total modeled using a 90% sequence similarity threshold on TCRα, TCRβ and peptide template selection, in order to prevent entries with high similarity to existing solved structures being more accurately modeled. There was no restriction on MHC template sequence similarity, as it was assumed that structural variability for all relevant MHC molecules was sufficiently described.

### Batch sampling experiment

For each peptide, binding and non-binding complexes were partitioned into batches of 1 binder and 5 non-binders, such that the batch contains complexes all with the same pMHC, for which one TCR is positive and the remaining TCRs negative. Given a batch, a scoring method is then tasked to assign a score to each of the 6 complexes, such that the positive binding complex is ranked as the highest scoring complex. Performance metrics are then computed in a per batch manner, evaluating how often binding complexes are correctly ranked. Here, we use a true-positive rank metric we term TPR that expresses how many non-binding complexes are scored higher than the binding complex:

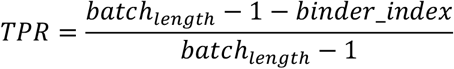

Additionally, we compute an additional metric we term batch accuracy. Here, a batch is considered correctly predicted if the binding complex is ranked as number one (bindex_idx == 0), and incorrectly predicted if not. The batch accuracy is then computed across all batches indicating the proportion of batches where the positive example was assigned the highest score (supplementary 2).

## Results

In order to effectively use structural models for predicting TCR-pMHC binding, we first set out to increase modeling quality by improving on the docking candidate selection step in AlphaFold. For this purpose, we developed a graph neural network based DockQ regressor, trained on a set of modeled TCR-pMHC complexes. Subsequently, we retrieved a large set of TCR sequences annotated with their peptide specificity from public databases and datasets, generated non-binding complexes by mismatching TCRs and pMHCs, and used our newly developed modeling pipeline to generate a set of binding and non-binding TCR-pMHC complex structural models. Using these structures, we then applied a range of deep learning based methods in order to classify binding and non-binding complexes.

### Structural diversity of AlphaFold-Multimer models

Initial experiments with modeling TCR-pMHC complexes using AlphaFold-Multimer (AF-M), revealed that for some benchmark dataset targets, many candidates would assume the same, incorrect configuration of CDR loops. This in turn led to loss of important native contacts of CDR3 loops, that would presumably hinder useful inference about the modeled structures and their immunological properties. We note that, we here use the benchmark dataset as a validation set for the development of the modeling pipeline, in order to quantify its capabilities for modeling the large set of sequences in the TCR specificity dataset. Inspired by Stein and Mchaourab 2022, we developed a set of tools to increase stochasticity in the AlphaFold-Multimer featurization pipeline and consequently reduce homogeneity of modeled candidates [22]. A set of candidate structures was then modeled for targets in the benchmark dataset, using a combination of different perturbation configurations and the effect of the perturbation was quantified w.r.t. structural diversity using the median intra-target pairwise backbone RMSD and w.r.t to modeling quality quantified by the DockQ metric, for details refer to materials and methods (figure 2).

**Figure 2:**
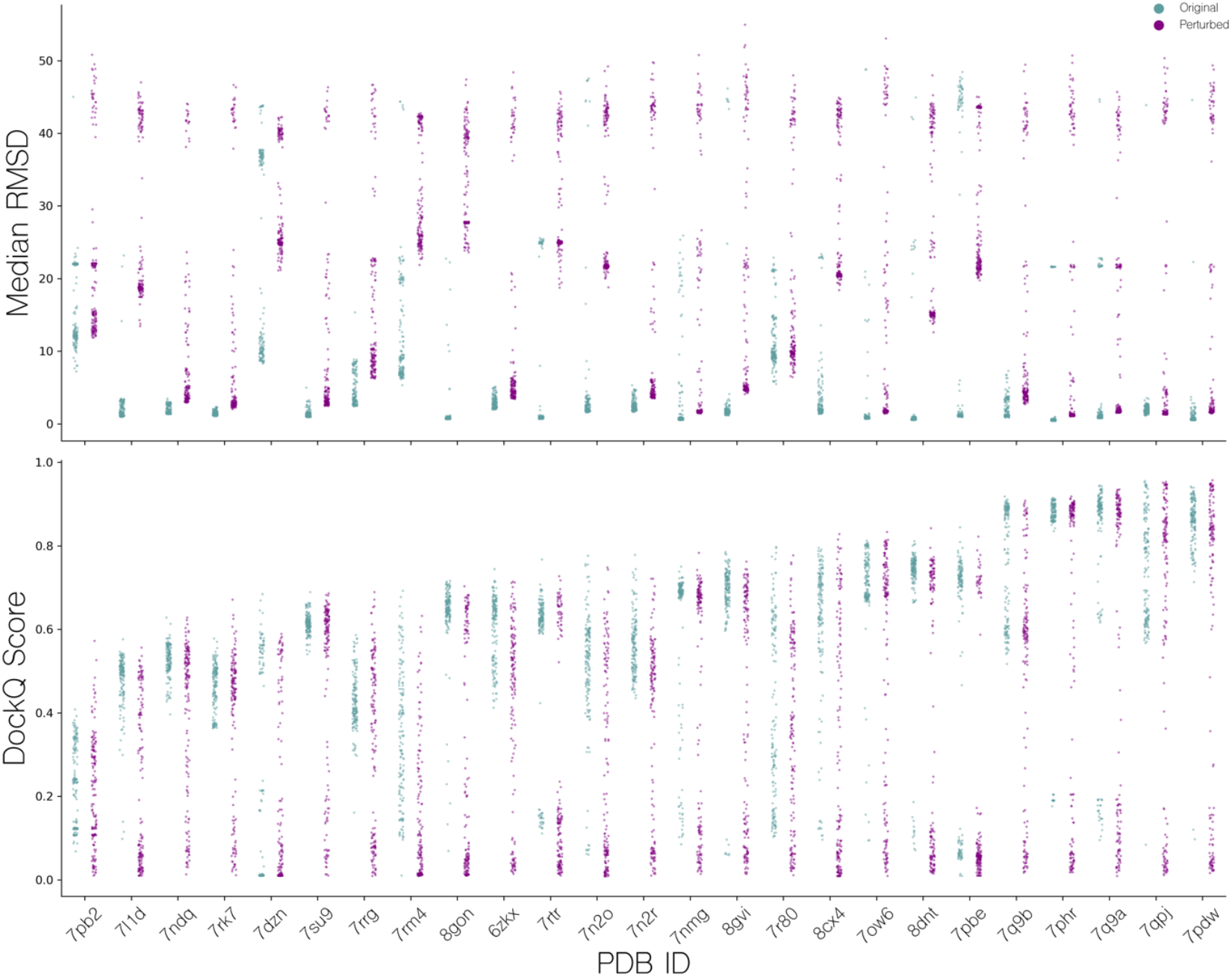
Distribution of quality scores for candidate structures modeled using AlphaFold-Multimer. Top: Median pairwise TCR RMSD (superimposed on pMHCs), for all candidates for a given target, before and after feature perturbation. Bottom: Distribution of DockQ scores for all candidates of a given target, before and after MSA and template feature perturbation.

When not applying MSA and template feature perturbation, the median of the per target median of median pairwise TCR RMSD is 1.47Å, indicating low structural diversity. This is especially pronounced for targets such as 7L1D, 7NDQ, 7RK7, where no or almost no candidate structures exceed a median pairwise RMSD of 4Å. Applying the feature perturbation method, the median of the per target median of median pairwise TCR RMSD values was increased to 6.12Å, showcasing a substantial increase in intra-target modeling diversity. However, this does not translate to an overall improvement in modeling accuracy, except for targets 7PB2, 7RK7 and 7RRG where we observe a max DockQ increase of approximately 0.1. Oppositely, other targets such as 7RM4, 7N2R and 7DZN have a comparative decrease in max DockQ. Additionally, we see a poorer density of high quality candidates which will make selection of high quality candidates harder, as the median of the median per target DockQ is brought down from 0.63 to 0.45. Based on these results, we opted not to use feature perturbation as a default feature of the pipeline and only use this approach for generating training data for the scoring function optimization.

### GVP-GNN regression performance and improvements over AlphaFold

Subsequently, we moved on the address the second challenge we encountered when modeling TCR-pMHC complexes using AF-M, namely that the confidence metric used internally in AF-M overestimates the quality for certain targets and consequently only correlates moderately with DockQ scores of candidates modeled for the benchmark dataset (see figure 3, upper panel). This figure demonstrates that a large number of complexes with AF-M quality scores above 0.7 have DockQ scores below 0.23 (corresponding to CAPRI threshold for incorrect structures [20]). We can further quantify this by calculating the cumulative proportion of incorrect structures (DockQ score less than 0.23) as a function of the annotated quality score (Figure 3 lower left panel). Doing this, we find that AF-M, at a predicted quality score of ∼0.7, has 2% of cumulative incorrect structures, within the set of ∼2200 top scoring complexes (Figure 3 lower right panel).

**Figure 3:**
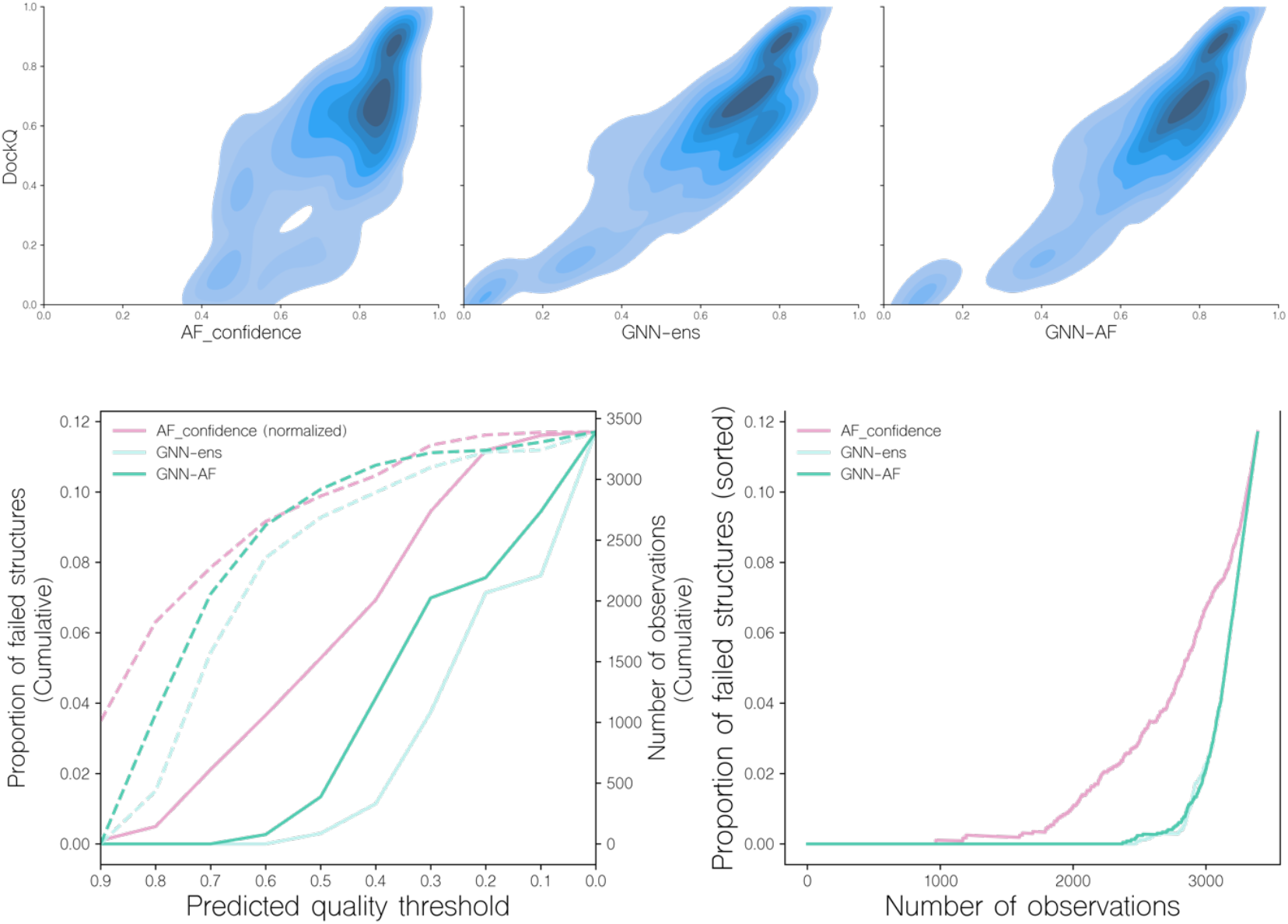
Distribution of DockQ scores for candidate structures ranked by different predicted quality scores. Top (a): Correlation of DockQ and AF_confidence, GNN-ens and GNN-AF for all candidates in the benchmark dataset. For more information on these scoring schemes, see the materials and methods section. Bottom left (b): The proportion of failed candidates in 10 bins of either AF_confidnece (normalized), GNN-ens, or GNN-AF score. Dotted lines indicate the number of observations in each bin. Following the CAPRI classification scheme, a candidate is considered as failed when DockQ < 0.23. Bottom right (c): The proportion of failed candidates as a function of the number of candidates, when sorted by either AF_confidence, GNN-ens or GNN-AF scores.

To address this issue, we developed a series of graph neural network based scoring methods to predict DockQ for a given input structure. Two GVP-GNN models were constructed, each trained in a 5 fold cross-validation setup, with data partitioned by reducing interpartitional joint TRA and TRB sequence similarity (for details see materials and methods). One was trained on the complete data set, and the other was trained only on docking poses with >0.5 DockQ (for details on the models and model training refer to materials and methods). The outputs from these two ensembles of 5 models were combined to predict DockQ for a given docking pose. We denote this score GNN-ens. Additionally, the outputs of these models were combined into a consensus score we denote GNN-AF, by computing the harmonic mean of the two GNN ensemble scores, AF_confidence and an additional AlphaFold score, AF_CDRpep_pLDDT (the mean of pLDDT scores associated with CDR123αβ and peptide residues, for more information on this score, refer to materials and methods).

Next, the performance of these quality assessment scores was evaluated on the validation and benchmark datasets in terms of different correlation metrics between the predicted quality and measured DocKQ and top1 Dock values (see Table 1). This analysis demonstrated an overall improved performance of the GNN-based models compared to AF-M. For instance, the overall Spearman’s rank correlation across all modeled structures improved from 0.681 to 0.824 (21%) for the GNN-ens model when compared to the AF-M confidence score. Notably, the correlation was greatly improved across the full DockQ range which in large parts resolves the issue of the AF-M confidence score overestimating the quality of poor quality candidates (Figure 3b, central panel). Analysing the rank correlation for candidates within each TCR-pMHC target also revealed a major increase of 52.02% (0.454 versus 0.297) increase in mean spearman correlation (refer to supplementary 1 for more information on the scoring of individual targets). Together this indicates that the GNN-ens method ranks candidates globally and locally better than the AlphaFold confidence metric. The performance gain compared to using AF-M score alone was further improved when considering the GNN-AF-ens model, merging the GNN ensemble with the AF-M confidence and AF-peptide-CDR pLDDT scores.

**Table 1:**
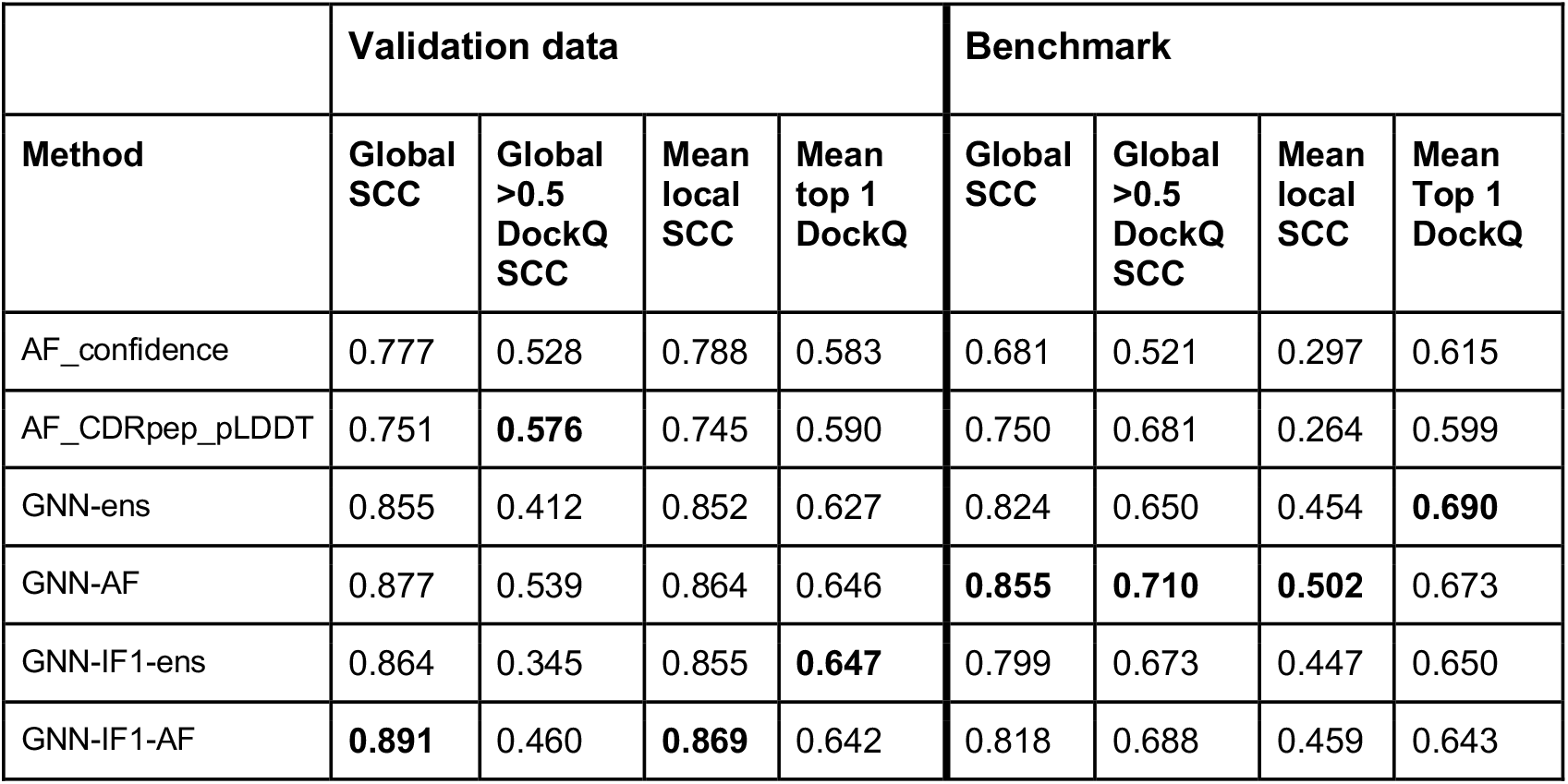
Scoring performance of GNN, GNN-IF1, AlphaFold and ensemble methods for the validation and benchmark data set. Performance metrics are, Global SCC: Spearman’s rank correlation coefficient (SCC) between predicted quality and DockQ, computed across all candidates for all targets, Global >0.5 DockQ SCC: SCC between predicted quality and DockQ, computed across all candidates for all targets with DockQ > 0.5, Mean local SCC: Mean per-target SCC between predicted quality and Do ckQ and Mean top 1 DockQ: Mean DockQ of the top 1 selection for each target. The GNN method is defined from the harmonic mean of the predicted DockQ score for two GVP-GNN ensembles; The GNN-AF score is computed by the harmonic mean of the two GVP-GNN ensemble scores, AF-M confidence and the peptide-CDR pLDDT score. The GNN-IF1 method is defined from the harmonic mean of the predicted DockQ score for two GVP-GNN-IF1 ensembles. The GNN-AF and GNN-IF1-AF models are defined as the harmonic mean of the two GVP-GNN (GVP-GNN-IF) ensemble scores, AF-M confidence and the AF peptide-CDR pLDDT scores. For details, see materials and methods.

|  | Validation data |  |  |  | Benchmark |  |  |  |
| --- | --- | --- | --- | --- | --- | --- | --- | --- |
| Method | Global SCC | Global >0.5 DockQ SCC | Mean local SCC | Mean top 1 DockQ | Global SCC | Global >0.5 DockQ SCC | Mean local SCC | Mean Top 1 DockQ |
| AF_confidence | 0.777 | 0.528 | 0.788 | 0.583 | 0.681 | 0.521 | 0.297 | 0.615 |
| AF_CDRpep_pLDDT | 0.751 | **0.576** | 0.745 | 0.590 | 0.750 | 0.681 | 0.264 | 0.599 |
| GNN-ens | 0.855 | 0.412 | 0.852 | 0.627 | 0.824 | 0.650 | 0.454 | **0.690** |
| GNN-AF | 0.877 | 0.539 | 0.864 | 0.646 | **0.855** | **0.710** | **0.502** | 0.673 |
| GNN-IF1-ens | 0.864 | 0.345 | 0.855 | **0.647** | 0.799 | 0.673 | 0.447 | 0.650 |
| GNN-IF1-AF | **0.891** | 0.460 | **0.869** | 0.642 | 0.818 | 0.688 | 0.459 | 0.643 |

Including the GNN-ens and GNN-AF-ens models in the analysis shown in Figures 3 further supports this gain in accuracy and demonstrates that at the score threshold where the proportion of failed structures reaches 2% for both methods correspond to ∼3100 complexes, an increase of 25% compared to the number of structures captured at the equivalent proportion of failed structures of AF-M.

The top 1 candidate selection is also notably improved (see Figure 4, and Table 1). Here, for almost all targets, a similar or better candidate was selected when using the quality score of the GNN or GNN-AF ensembles resulting in an increase of 12% (0.615 to 0.69) and 9.43% (0.615 to 0.673), respectively, in the mean DockQ of top 1 selections over the different targets in the benchmark data set compared to AF-M. Classifying the top 1 selected models according to the CAPRI quality categories, the GNN based methods further completely avoids selection of “Incorrect” candidates (Figure 4). Further, using the GNN-AF consensus score, we observe an increase in the proportion of “Medium” (0.49 <= DockQ < 0.80) and “Acceptable” (0.23 <= DockQ < 0.49) quality structures.

**Figure 4:**
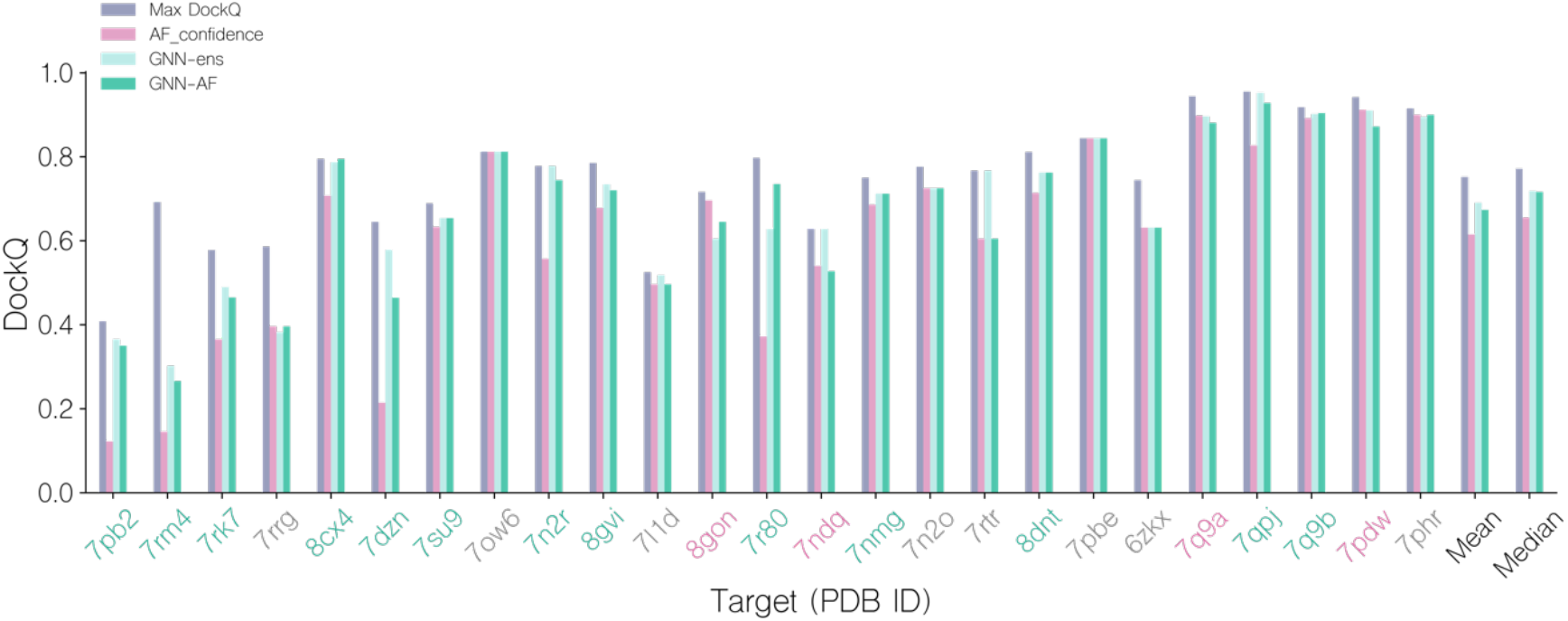
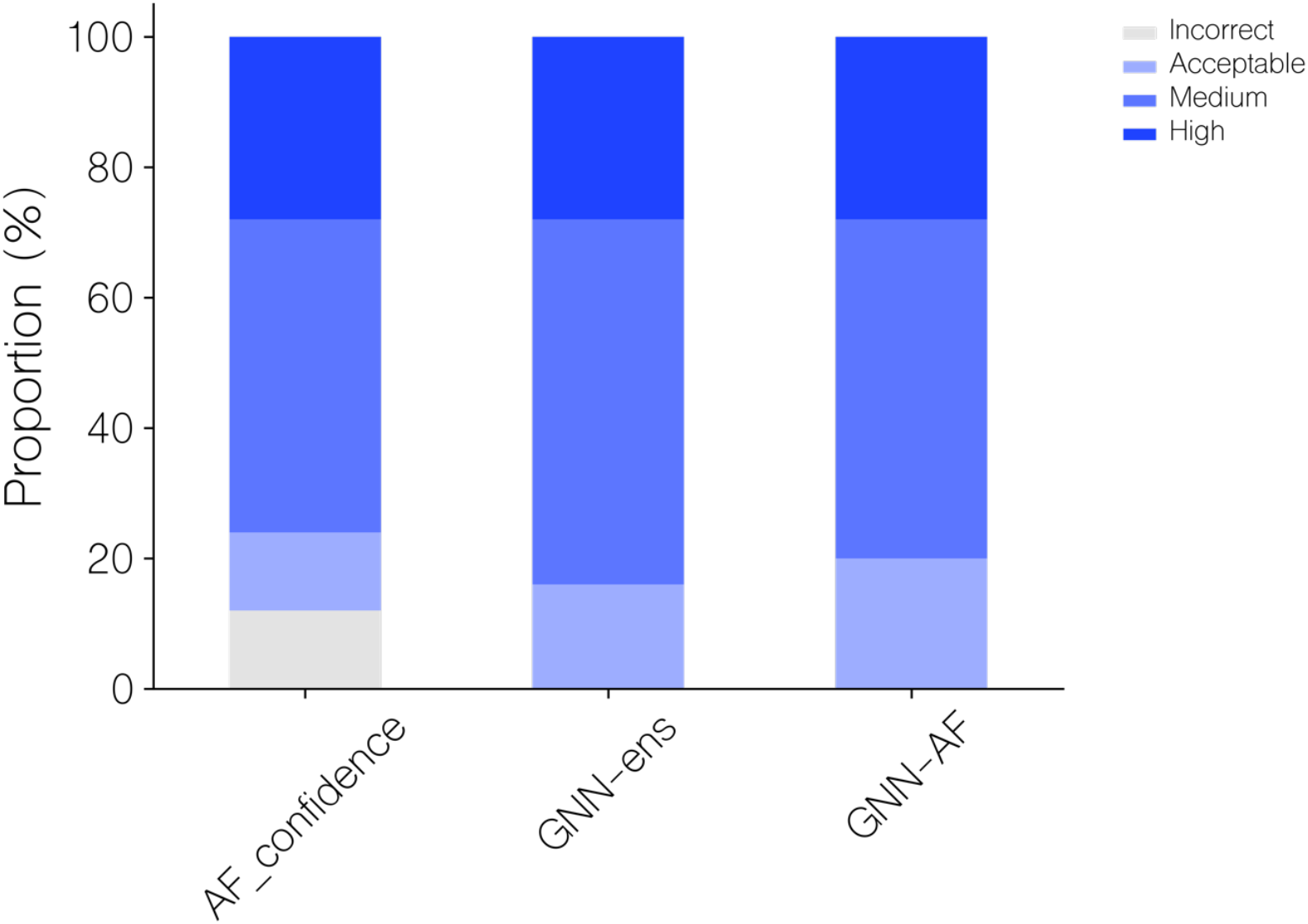
Performance evaluation of AF-M, GNN and GNN-AF quality prediction methods for top1 candidate selection. Top: DockQ of top 1 selections for each target, using AF_confidence, GNN-ens or GNN-AF scoring. DockQ of the highest quality candidate is also shown for each target. Red PDB ID label: AF_confidence results in a better selection, gray: identical selection, green: GVP-GNN based scoring results in a better selection. Targets are sorted by their top 1 GNN-AF score. Bottom: Proportions of CAPRI quality categories for top 1 selections for the three methods.

Based on these results, we opt to use the GNN-AF score for the subsequent analyses.

### Prediction of TCR-pMHC binding

Next, we turned to the challenge of predicting the correct TCR binding to a given pMHC target. Given the capabilities of the GNN-AF score for ranking docking poses, we hypothesized that it could also be used to separate cognate TCR-pMHC complexes from complexes with swapped incorrect pairings. The intuition being that swapped complexes to a lesser degree would resemble “real” binding complexes, and therefore would tend to be scored lower by quality evaluation metrics.

To investigate this, a TCR specificity dataset consisting of binding and swapped TCR and pMHC was created as described in the materials and methods section. Briefly, the dataset was generated by downsampling the TCR specificity dataset created in Jensen (2024). The downsampling was made to ensure a more even distribution in the number of complexes for each peptide, and to allow for sampling of a large number of structural models for each complex, which would otherwise not be possible for the complete dataset due to computational complexity. The resulting data set consists of 2,945 binding complexes each matched with 5 swapped negative (i.e. the pMHC matched with TCRs positive to other pMHCs) complexes, resulting in 14,725 = 5 * 2,945 negatives giving a total of 17,670 examples spanning 26 peptides (for more details, see materials and methods).

The different methods were next evaluated in a batch setup, where each batch contained one positive and 5 swapped negative complexes. Structures for each complex were modeled as described in the materials and methods section and the top 1 candidate for each complex selected using a given scoring approach. Next, the complexes within the batch were scored and a metric termed TPR (for details see materials and methods) was computed, which quantifies how many of the negative (swapped) complexes in the batch were scored higher than the positive example. A TPR of 1.0 thus indicates that the positive example was assigned the highest score in the batch. We then sorted batches by the highest intra-batch score predicted by a scoring method in order to compute a cumulative TPR curve for all batches (Figure 5a). Here, we further included a variant of the GNN method trained including the ESM-IF1 inverse folding embedding as input for each residue in the network. This was done to investigate if such representations could serve to boost the method’s ability to discriminate the binding interface between true and swapped complexes. When evaluated in terms of top 1 candidate selection and correlation between predicted quality and DockQ values, this method demonstrated a comparable performance to that of the GNN method, both alone and in combination with AF-M (see Table 1).

**Figure 5:**
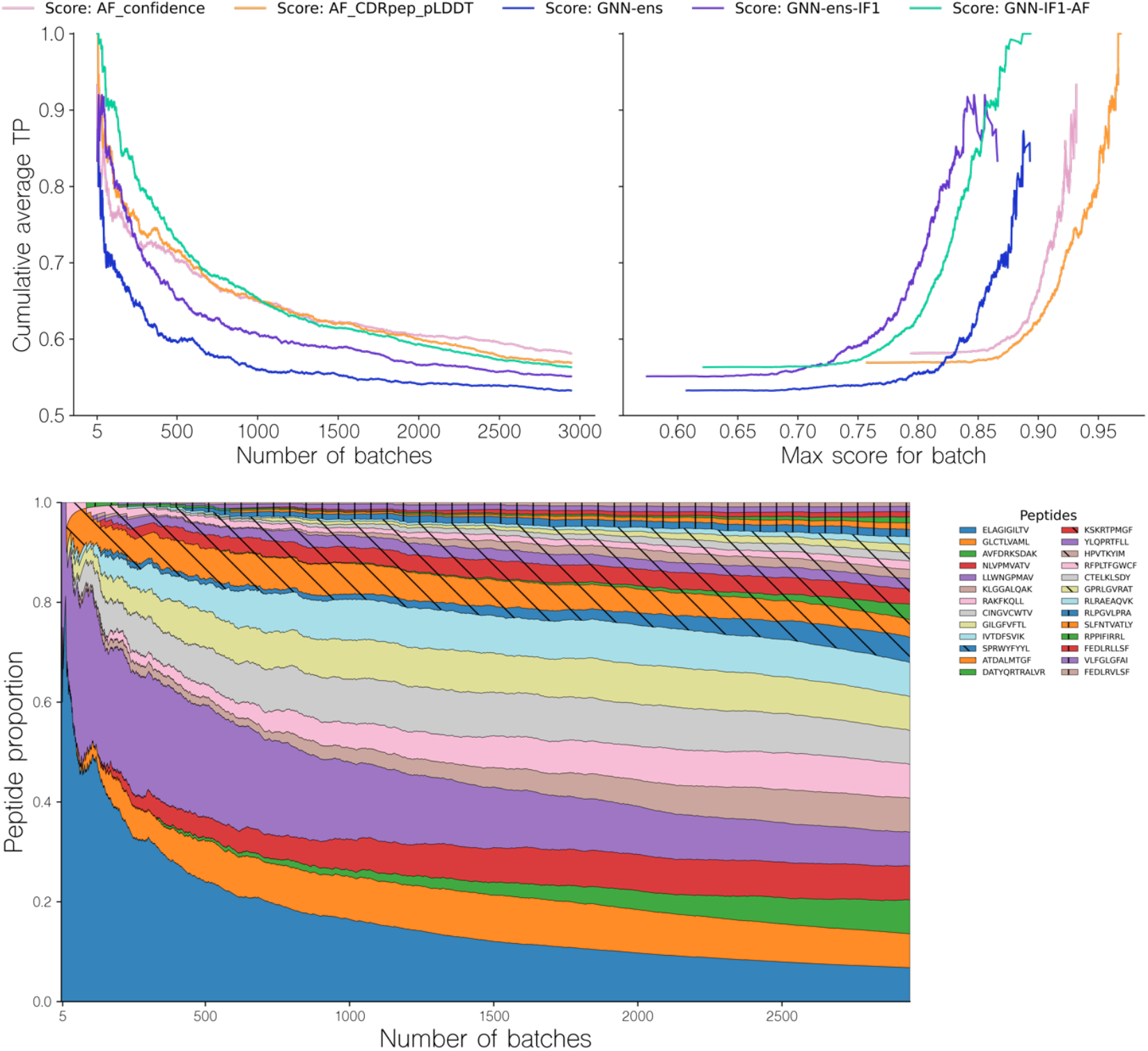
Performance evaluation of the different methods in the batch evaluation. Upper panel Left (a): Cumulative TPR curve for batches sorted by descending maximal intra-batch quality score for a given method. The TPR value for a given batch is computed from the ranking of the binding complex within each batch, based on their predicted quality, with a value of 1 corresponding to a top 1 rank. Upper panel Right (b): Cumulative TPR curve for batches as a function of the maximal intra-batch quality score. The different methods are, AF_confidence: AlphaFold-M confidence, AF_CDRpep_pLDDT: Mean pLDDT of CDR123 and peptide residues, GNN-ens: GVP-GNN ensemble, GNN-ens-IF1: GVP-GNN ensemble trained including ESM-IF1 structural embeddings.

GNN-IF1-AF: Harmonic mean of GNN-ens-IF1, AF_confidence and AF_CDRpep_pLDDT. For more information on the GVP-GNN ensembles, see text and materials and methods. Bottom (c): Proportion of observed peptides in a range of batches sorted according to their max intra-batch score. Note that to avoid showing averages over small numbers, the first data point shown in all plots corresponds to the top 5 batches.

Figure 5 demonstrates that methods combining GNN and AlphaFold-M scoring metrics tend to have a greater cumulative TPR curve AUC, specifically for the first 1000 batches, where GNN-IF1-AF achieves an AUC of 0.76, compared to AUCs of 0.71, 0.72 and 0.69 for AF_confidence, CDRpep_pLDDT and GNN-ens-IF1 respectively. By way of example, the GNN-IF1-AF ensemble maintains a cumulative TPR above 0.8 (corresponding to an average rank of top 2 within the batch size 6) for the top ∼300 (∼10.2%) batches. This suggests that when a batch contains sufficiently accurate structural models, this consensus method can accurately separate binding complexes from non-binders. In contrast, the individual scoring methods, AF_confidence, AF-CDRpep_pLDDT, GNN-ens-IF1 and particularly GNN-ens archives this high accuracy for a much smaller set of batches. Here, the drop below 0.8 in cumulative TPR occurs already at 60 (2%) and 110 (3.7%) batches for AF_confidence and AF-CDRpep_pLDDT respectively. Notably, the GNN-ens score performed worse in selecting binding complexes compared to the other methods, despite its superior performance in ranking and selecting high modeling quality docking poses (Table1). It is not clear what is driving this drop in performance. Therefore, from these results, it is evident that combining the AlphaFold and GNN-ens-IF1 scoring methods into a consensus score, yields superior predictive power for selecting binding TCR-pMHC complexes. Further examining the TPR as a function of the maximal batch scores of the various scoring methods allows us to identify a threshold value at which we can expect accurate ranking within a batch (figure 5b). Focusing on the consensus GNN-IF1-AF score, we see that the average accumulated TPR of 0.8 corresponds to a predicted quality score of approximately 0.825.

Plotting the distribution of peptides observed for each of batches sorted by the GNN-AF-quality score, we observe that high scoring batches are primarily observed for the peptides LLW and ELA (figure 5c). That is of the top 100 batches, 79.2% are for the peptides LLW and ELA. This distribution is however more spread out within the top 300 batches (corresponding to the point where the cumulative TPR falls below 0.8). Here, a total of 7 peptides each contribute more than 5% to the distribution.

In conclusion, these results suggest that the proposed quality assessment scoring scheme can accurately separate correct from wrongly matched TCR-pMHC paired in cases where the predicted structural accuracy is high. The results however also indicate that is the case in relatively few instances.

### Structural inaccuracies may obfuscate binder signal

Given these results and the observation that few TCR-pMHC complexes were modeled with sufficiently high quality to allow for reliable target assessment, we hypothesized that the structure pipeline was unable to produce structural models with sufficiently accurately docked TCRs. To further investigate this, we first recreated the binding classification for models from the structure modeling benchmark dataset (for details refer to materials and methods). However, given the low number of structures, we opted here not to use homology reduction on this set, giving us 38 structures as opposed to the 25 as described above. For each of the 38 complexes, now denoted as the “binding” complexes, 5 non-binding complexes were generated by swapping the TCR with TCRs randomly selected from other complexes. To avoid sampling potential cross-reactive TCRs as negatives, for a given complex TCRs were only sampled from complexes with a peptide Levenshtein distance to the original complex of at least 3. This resulted in a dataset of 228 complexes. Following this, top 1 candidate structures were selected for each complex using the various scoring methods, and the batch sampling experiment was repeated (Figure 6). For this dataset, the AF confidence metric struggles to correctly rank binding complexes, achieving average cumulative TPR of only 0.5 for the first 20 batches, equivalent to random ranking. Notably, the AF_CDRpep_pLDDT score achieves the highest TPR AUC out of all methods on this dataset, maintaining an average cumulative TPR above 0.6 for the first 10 batches.

**Figure 6:**
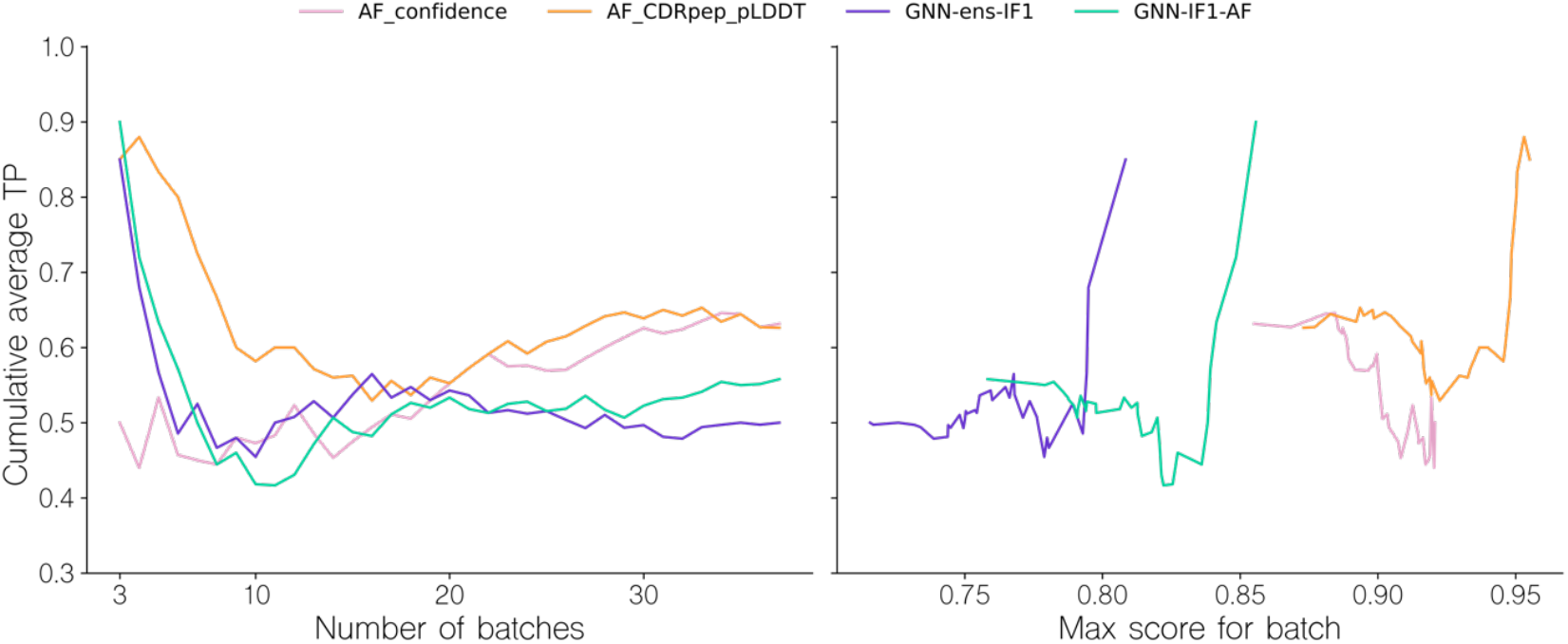
Batch sampling TPR curves for benchmark dataset models. Left (a): Cumulative TPR curve for batches sorted by their TPR. The TPR values are computed by how accurately binding and swapped complexes are ranked within each batch, based on their predicted quality. A TPR closer to 1.0 indicates more correct ranking within batches. Right (b): Cumulative TPR curve for batches sorted by the max intra-batch quality score. Note that to avoid showing averages over small numbers, the first data point shown in all plots corresponds to the top 3 batches.

The GNN based methods, GNN-ens-IF1 and GNN-IF1-AF archive high TPR values only for the first 5 (13%) batches, but subsequently quickly fail to produce accurate rankings. However, the drop in cumulative TPR happens around the same quality_score threshold of 0.825 as observed in Figure 5, corroborating the idea that we can use the maximal intra-batch quality score to determine which batches can be accurately ranked. These results, thus overall align with the finding from the larger-scale NetTCR binding data set of Figure 5. This means, we potentially from this structural data set can investigate properties related to modeling quality, and point to sources for their low accuracy.

For this, we in Figure 7 first display DockQ and the predicted GNN-IF1-AF quality scores of the complexes generated for the positive TCR-pMHC examples as a function of the mini-batch TPR. Here, we can observe significant correlation coefficients between modeling quality measures and classification success (r=0.65, p-value =9.8 10^-6, two-tailed t-test test). Focusing on the aggregate DockQ quality measure, we observe that moderately high model quality (⪆0.6 DockQ) is a needed but not sufficient prerequisite for achieving batch TPR values over 0.5. Further, notably, only batches with very high quality positives (⪆0.8 DockQ) are all ranked very accurately, with most batches achieving a TPR of 1.0. This is corroborated by the GNN-IF1-AF score, which is shown to be more strongly correlated with TPR (r=0.88, p-value =1.7 10^’13, two-tailed t-test test),with batches with a positive complex with a score above 0.8 all archiving a TPR value 0.8 or above. These results show that structural modeling quality clearly influences TCR-pMHC binding classification success. Particularly, that high quality models (⪆0.8 DockQ and ⪆0.8 GNN-IF1-AF score) are required for accurate classification. To further this analysis, we computed CDR3ab RMSDs after superimposing the models onto the pMHC and examined their relationships with predicted quality and TPR (Figure 8).

**Figure 7:**
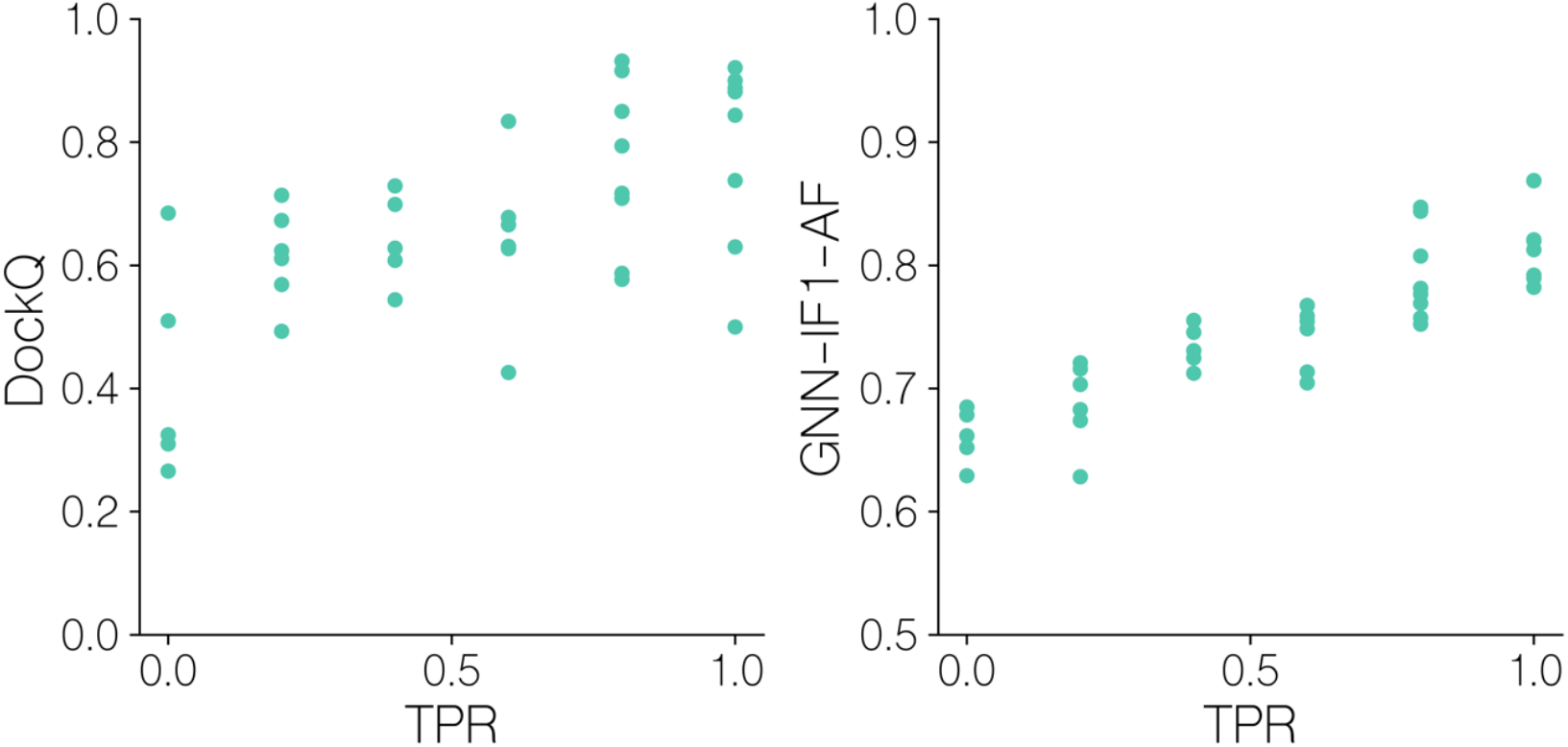
Batch sampling TPR values versus quality metrics for benchmark dataset models. DockQ and GNN-IF1-AF scores versus mini-batch ranking TPR for models of solved structures. GNN-IF1_AF: combined GVP-GNN-IF1 ensemble, AF-M confidence and peptide-CDR pLDDT scores (see materials and methods). A TPR of 1.0 indicates that in the given batch, the positive example was assigned the highest GNN-IF1-AF score.

**Figure 8:**
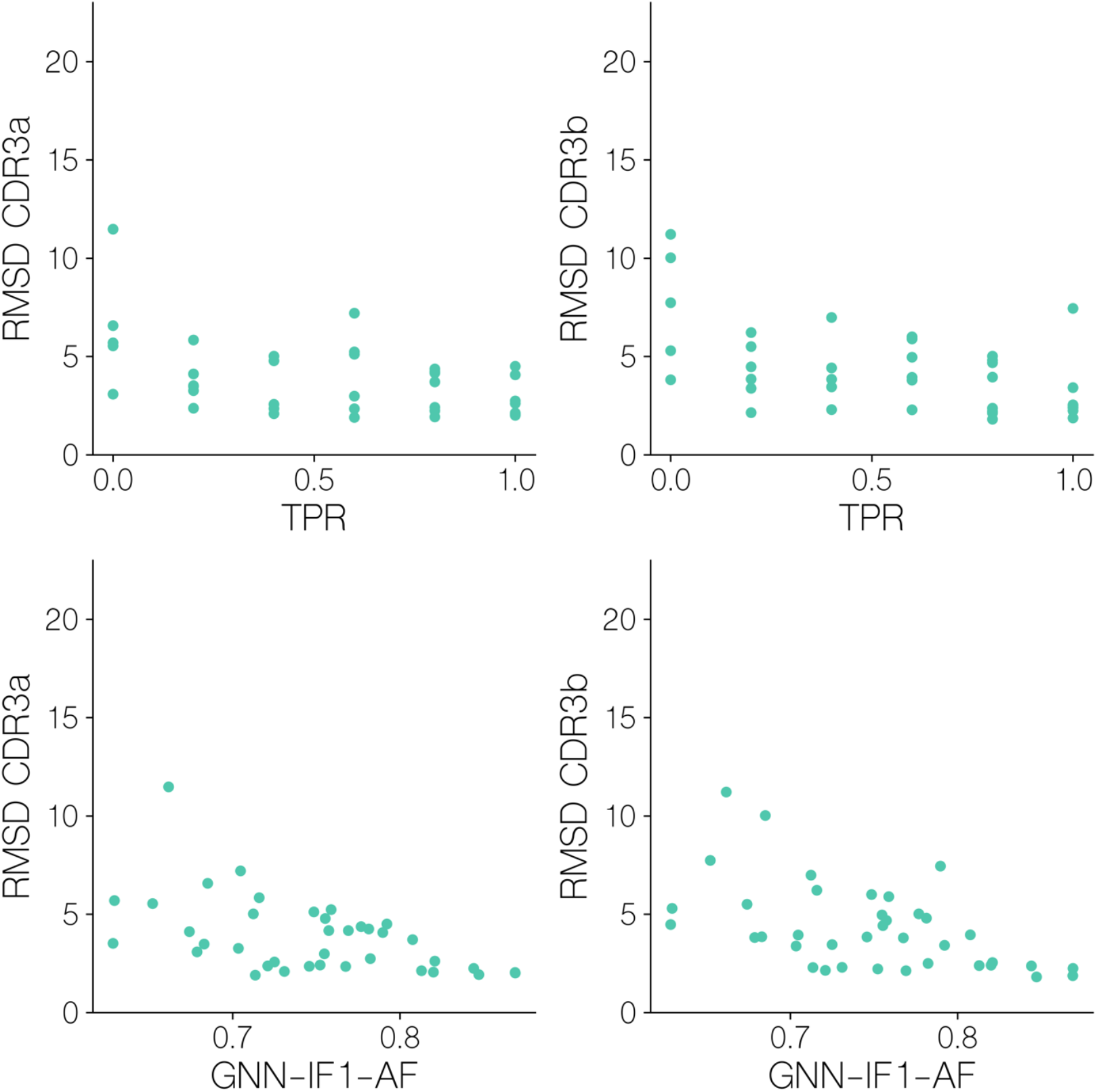
CDR3 atom coordinates RMSD values versus scoring metrics. CDR3ab backbone/sidechain RMSD after superimposition on pMHC CA atoms for benchmark set targets, versus their GNN-IF1-AF score and mini batch TPR.

Here, we observe that for both CDR3a and CDR3b backbone/sidechain atoms, an RMSD of <2.5Å is required in order to reach the aforementioned GNN-IF1-AF score threshold of ∼0.8 that predicts classification success, that we see very few structural models reach. We see that mini-batch TPR is significantly correlated with both CDR3a and CDR3b RMSD (r=-0.462, p-value=3.52 10^-3, r=-0.529, p-value=6.43 10^-4, two-tailed t-test), showing that docking accuracy significantly influences how accurately we can assess binding and non-binding complexes. Particularly for CDR3b loop, with the exception of an outlier, all batches with a TPR of 1 have an RMSD of <2.5Å. This again, suggests a modeling quality threshold that is difficult to attain when also considering findings from the modeling quality benchmark (figure 4) and the observed classification performance (figure 5).

### Predictive performance on unseen peptides in IMMREP23

In order to further validate the predictive performance of the GNN-IF1-AF score, we evaluated the model on a subset of the IMMREP23 TCR specificity prediction benchmark dataset [9].Specifically, we focused on data for the previously unseen peptides SALPTNADLY, TSDACMMTMY and FTDALGIDEY. We modeled the 246 TCR-pMHC complexes from this subset using the modeling pipeline described here and scored the resulting models using a set of different scoring methods including AF-M, GNN and the GNN-IF1-AF ensemble scores. However, for this dataset we opted to only model 10 candidates per complex and without any restrictions on template selection in order to mimic a more realistic use case of the modeling and scoring pipeline. We then selected the top 1 scoring complex, using each of the scoring methods, and used this score to compute an AUC 0.1 value for each peptide (Figure 9).

**Figure 9:**
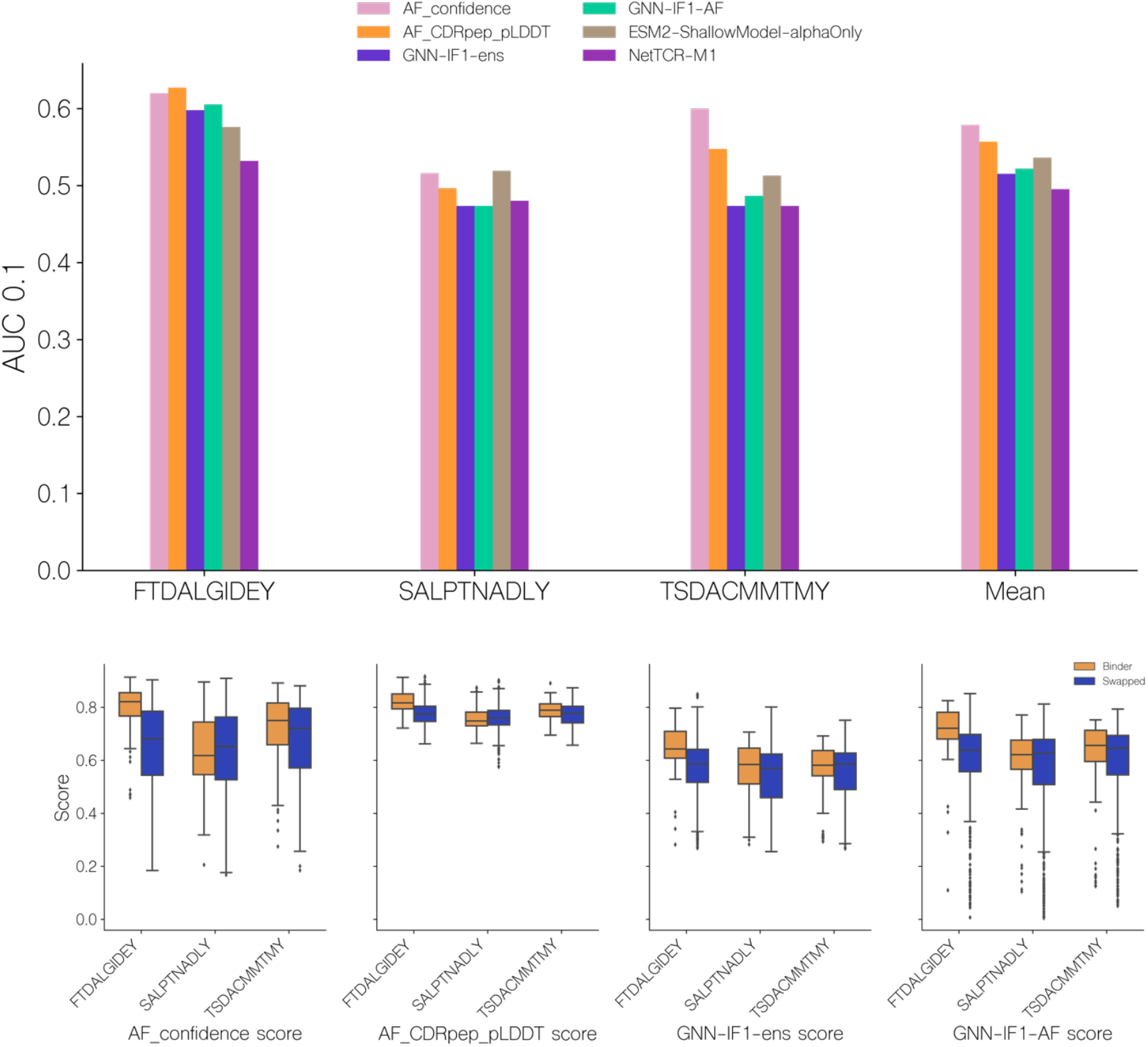
Predictive performance on a subset of the IMMREP23. Top (a): Predictive performance of quality scoring methods on data for the unseen peptides in the IMMREP23 benchmark dataset. The AUC 0.1 is computed from the predicted quality score, obtained from the top 1 score of each scoring method. , AF_confidence: AlphaFold-M confidence, AF_CDRpep_pLDDT: Mean pLDDT of CDR123 and peptide residues, GNN-ens: GVP-GNN ensemble, GNN-ens-IF1: GVP-GNN ensemble trained including ESM-IF1 structural embeddings. GNN-IF1-AF: Harmonic mean of GNN-ens-IF1, AF_confidence and AF_CDRpep_pLDDT. For more information on the GVP-GNN ensembles, see text and materials and methods. ESM2-ShallowModel: A 3-layer multilayer perceptron train on TCR and peptide protein sequence embeddings from ESM2. NetTCR-M1: 1 dimensional convolutional neural network trained on BLOSUM50 embeddings of TCR and peptide sequences [27]. Bottom (b): Distribution of predicted quality scores for each of the unseen IMMREP23 benchmark peptides.

On this benchmark dataset, we find that the AlphaFold-M scores, AF_confidence and AF_CDRpep_pLDDT achieved a slightly higher AUC0.1 than the GNN based scores. This is the case for all three peptides, however mostly pronounced for TSD, where the AF_confidence and AF_CDRpep_pLDDT scores achieve AUC0.1 values of 0.60 and 0.548 respectively, while the GNN based methods, GNN-IF1-ens and GNN-IF1-AF, achieve values of 0.474 and 0.487 respectively. For the remaining two peptides, all methods perform similarly. On SAL, performance is poor across all methods, with AUCs around 0.5, while on FTD, they all achieve AUCs of approximately 0.6 indicating better than random classification. Plotting the distribution of scores predicted by the different methods for each peptide (Figure 9b), we find all methods share higher scoring values for the FTD peptides, which the methods can more accurately predict binding for. For the GNN-IF1-AF method in particular, only FTD has complexes with scores >0.8, where 12% and 2% of binders and non-binders respectively surpass this threshold. The proportion of complexes with a score >0.7, for each peptide for binders and non-binders are 61% and 24%, 15% and 19% and 34% and 22% for peptides FTD, SAL and TSD respectively. Thus, we see an enrichment in higher quality structural models for binders, for 2 out of 3 peptides. These observations corroborate our hypothesis, that only in cases with high quality structural models, we can use these to predict binding. We also here compared the performance of these scoring methods to the performance of two models that were entered in the IMMREP23 benchmark. Both of these models use amino acid sequence data to generate predictions, where one is a multilayer perceptron taking ESM-2 protein language model embeddings as input and the other is a convolutional neural network taking BLOSUM50 embeddings as input [27], [28]. Here, we find that all methods based on structural data outperform the sequence based models on FTD, and the AlphaFold-M scores for TSD. While the methods utilizing GNN ensembles in this particular benchmark only achieved similar performance to the sequence based methods, these results demonstrate that structural data can assists in predicting TCR-pMHC binding for novel peptides.

### NetTCR-struc web server

We have made our GNN-AF and GNN-IF1-AF methods available in a GitHub repository for use in docking scoring and TCR-pMHC binding prediction at https://github.com/mnielLab/NetTCR-struc.

## Discussion

In this work, we made efforts to improve structural modeling accuracy of TCR-pMHC class I complexes and evaluated the use of structural data for predicting TCR-pMHC binding. For these tasks, we evaluated a range of methods, including AlphaFold-Multimer’s internal scoring metrics as well graph neural network based methods trained on structural models of TCR-pMHC complexes.

## Structural diversity and modeling accuracy

Initially, we observed that AF-M produced structurally homogeneous docking candidates, often misrepresenting the CDR loop configurations. Using MSA feature perturbation, we demonstrated that we could generally improve modeling diversity, however this did not lead to any meaningful improvements in overall modeling quality. However, given its mixed effects on modeling accuracy, we opted to use this approach selectively for training a scoring function rather than as a default setting.

A major challenge in model selection was the overestimation of model quality by AF-M’s internal confidence metric. Our analysis revealed only a moderate correlation between AF-M confidence scores and DockQ scores, leading to incorrect assessment of low-quality models. To improve this, we developed a series of GNN-based scoring methods trained to predict DockQ scores. Our best-performing model, GNN-AF, combined GNN predictions with AF-M confidence metrics, yielding a 21% increase in Spearman’s correlation (from 0.681 to 0.855) with DockQ, significantly improving ranking accuracy for docking candidates.

Touching upon DockQ as a metric for evaluating TCR-pMHC modeling quality, we noted that the DockQ values reported for the complexes in the benchmark set appeared to be generally high with a mean of 0.673 DockQ for the top 1 selections. However, due to the highly conserved nature of TCR-pMHC docking geometry these numbers are in fact an exaggeration of the actual modeling skill of AlphaFold-M. Thus, while some targets such as 7PHR, 7Q9B, 7PWD, 7Q9A and 7QPJ that are modeled with near-native quality likely do indeed capture most pMHC and TCR interactions, models for the remaining targets, likely fail to capture important interactions in the docking interfaces, despite their generally high DockQ values.

### Prediction of TCR-pMHC binding

Using the methods developed in the docking pose quality scoring, we explored the potential for using modeling quality as a score for TCR-pMHC binding prediction. Here, our results demonstrated that integrating GNN-AF with inverse folding embeddings (GNN-IF1-AF) further improved discrimination of binding versus non-binding complexes, particularly when high-quality structural models were available. However, while the GNN-IF1-AF ensemble outperformed AF-M scoring metrics, it (along with all other methods) only maintained a high performance for a small proportion of analysed data. This therefore underlined that the structural pipeline in general struggled to generate sufficiently high-quality TCR-pMHC models suitable binding prediction. This was further elucidated when evaluating binding classification performance on the solved structures docking pose ranking benchmark dataset. Here, the AF confidence metric failed completely to effectively rank binding complexes, whereas the GNN-IF1-AF method performed better for a small subset of high quality structures but otherwise failed. This again suggests that structural modeling quality is a key determinant of ranking performance.

While conducting this research, AlphaFold-3 was released which demonstrates significant improvements in modeling quality, particularly for antibodies. Due to the related nature of TCRs and antibodies, we might expect this version to also showcase improvements in modeling for TCR-pMHC complexes over AlphaFold-2.3. However, since the docking scoring approach we have developed here is orthogonal to the AlphaFold scoring scheme, we still expect the approach of applying (a potentially refined) TCR-pHLA specific quality scoring model would help improve ranking of structural models created also using AlphaFold-3. The same argument holds when it comes to our conclusions on the limitations of structural modeling and how it affects TCR-pMHC binding. While the newest structural modeling approaches might improve on this, we still expect the challenge to remain for parts of the TCR-pMHC binding space. To access and characterize this, we recommend further investigations are conducted along the lines of the work described here.

In conclusion, we have here presented a pipeline incorporating inverse-folding embeddings and GNN-based scoring for refined quality assessment of TCR-pMHC structures. This approach was found to significantly improve structural ranking and binding prediction, and further provided insights into how structural modeling quality affects binding prediction performance. However, our findings also elucidated key limitations in current structural modeling methods. The release of AlphaFold-3 may offer improvements in modeling quality, thereby potentially addressing some of the observed shortcomings. However, our results suggest that domain specific, novel quality scoring approaches like those present here will remain useful for improving on their corresponding structural modeling tasks. Continued work on this topic will allow limitations both in terms of the structural modeling and scoring accuracies to be identified and characterized, enabling new pathways for refining docking scoring approaches and subsequently improving the accuracy and reliability of TCR-pMHC binding predictions.

