## Supplementary figures for "NetTCR-struc, a structure driven approach for prediction of TCR-pMHC interactions"

### Supplementary material

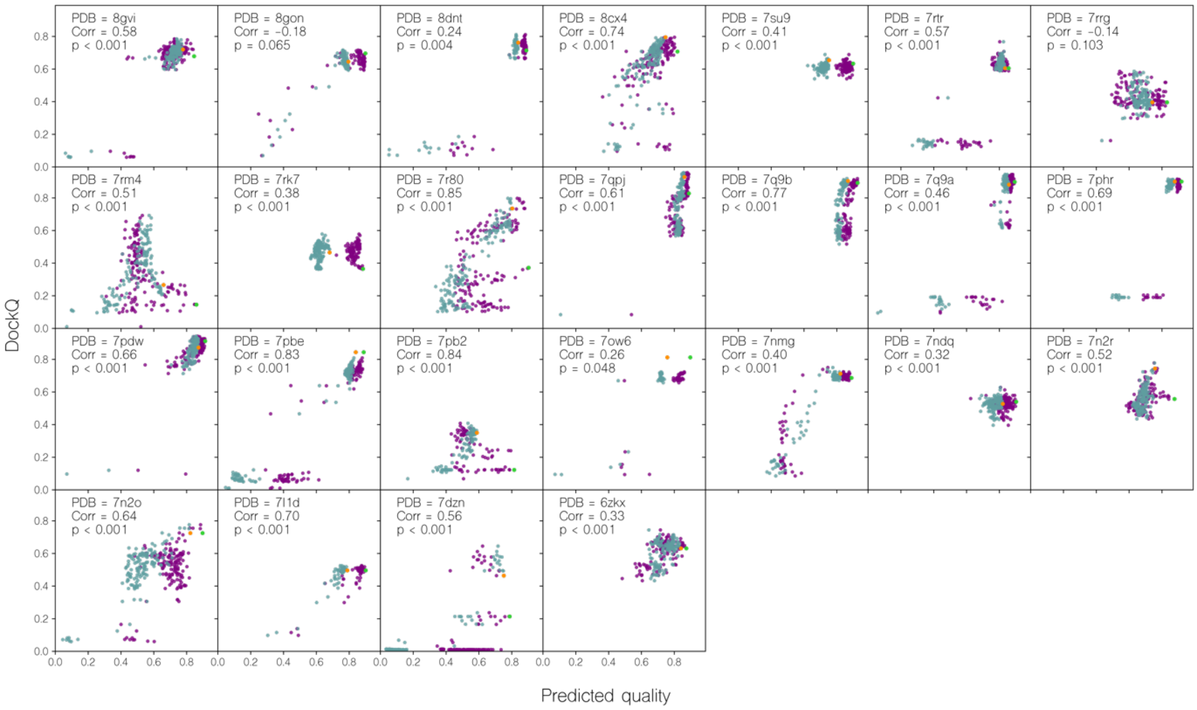


**Supplementary 1: Per benchmark target correlation of DockQ and predicted quality.** Green: GNN-AF. Purple: AF_confidence. Orange and green dots indicate the highest confidence for each target, i.e. the candidate that would be selected as top 1 for GNN-AF and AF_confidence respectively. The Spearman correlation coefficient for GNN-AF and DockQ is shown in each subplot.

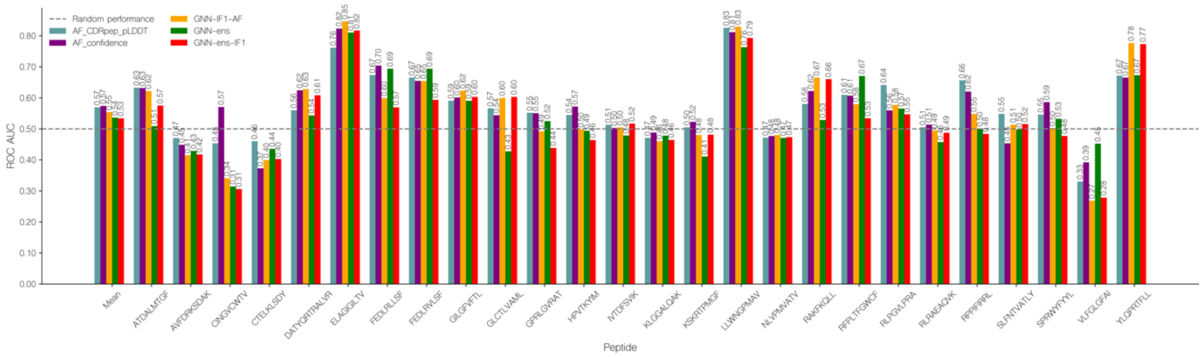


**Supplementary 2: TCR-pMHC binding classification ROC AUC for different scoring methods.** The different methods are, AF_confidence: AlphaFold-M confidence, AF_CDRpep_pLDDT: Mean pLDDT of CDR123 and peptide residues, GNN-ens: GVP-GNN ensemble, GNN-ens-IF1: GVP-GNN ensemble trained including ESM-IF1 structural embeddings. GNN-IF1-AF: Harmonic mean of GNN-ens-IF1, AF_confidence and AF_CDRpep_pLDDT. For more information on the GVP-GNN ensembles, see text and materials and methods.

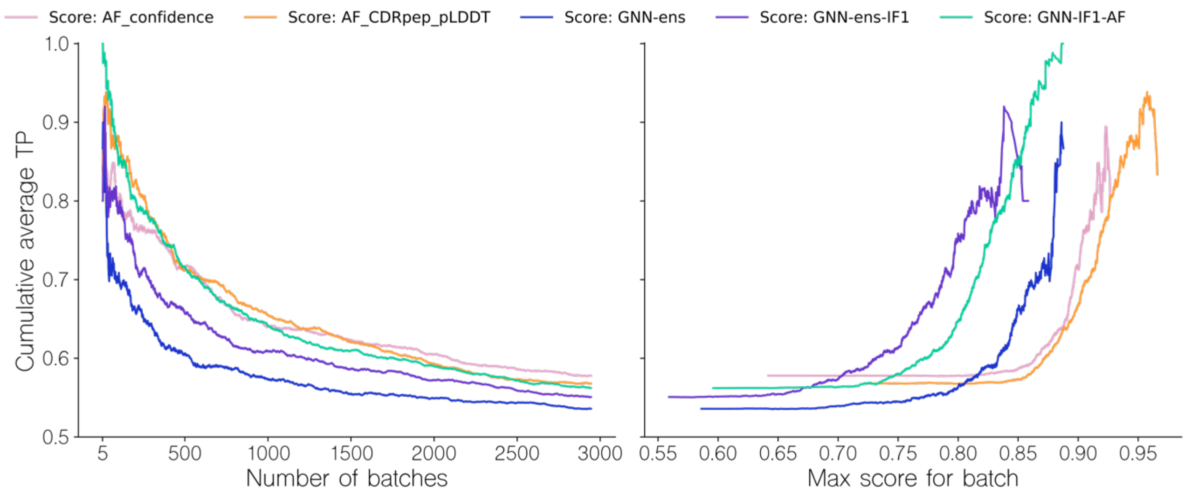


**Supplementary 3: Performance evaluation of the different methods in the batch evaluation when selecting top 1 based on GNN-AF.** Left (a): Cumulative TPR curve for batches sorted by descending maximal intra-batch quality score for a given method. The TPR value for a given batch is computed from the ranking of the binding complex within each batch, based on their predicted quality, with a value of 1 corresponding to a top 1 rank. Right (b): Cumulative TPR curve for batches as a function of the maximal intra-batch quality score. In both plots, docking candidate selection was performed using only the GNN-AF score. The different methods are, AF_confidence: AlphaFold-M confidence, AF_CDRpep_pLDDT: Mean pLDDT of CDR123 and peptide residues, GNN-ens: GVP-GNN ensemble, GNN-ens-IF1: GVP-GNN ensemble trained including ESM-IF1 structural embeddings. GNN-IF1-AF: Harmonic mean of GNN-ens-IF1, AF_confidence and AF_CDRpep_pLDDT. For more information on the GVP-GNN ensembles, see text and materials and methods.
